# Long-term potentiation in neurogliaform cells modulates excitation-inhibition balance in the temporoammonic pathway

**DOI:** 10.1101/531822

**Authors:** Marion S. Mercier, Vincent Magloire, Jonathan H. Cornford, Dimitri M. Kullmann

**Author notes:** Mila, Montreal, QC, H2S 3H1, Canada.

## Abstract

Apical dendrites of pyramidal neurons integrate information from higher-order cortex and thalamus, and gate signaling and plasticity at proximal synapses. In the hippocampus, neurogliaform cells and other interneurons located within stratum lacunosum-moleculare mediate powerful inhibition of CA1 pyramidal neuron distal dendrites. Is the recruitment of such inhibition itself subject to use-dependent plasticity, and if so, what induction rules apply? Here we show that interneurons in mouse stratum lacunosum-moleculare exhibit Hebbian NMDA receptor-dependent long-term potentiation (LTP). Such plasticity can be induced by selective optogenetic stimulation of afferent fibers in the temporoammonic pathway from the entorhinal cortex, but not by equivalent stimulation of afferents from the thalamic nucleus reuniens. We further show that theta-burst patterns of afferent firing induces LTP in neurogliaform interneurons identified using neuron-derived neurotrophic factor (Ndnf)-Cre mice. Theta-burst activity of entorhinal cortex afferents led to an increase in disynaptic feed-forward inhibition, but not monosynaptic excitation, of CA1 pyramidal neurons. Activity-dependent synaptic plasticity of neurogliaform cells in stratum lacunosum-moleculare thus alters the excitation-inhibition balance at entorhinal cortex inputs to the apical dendrites of pyramidal neurons, implying a dynamic role for these interneurons in gating CA1 dendritic computations.

**Significance statement:** Electrogenic phenomena in distal dendrites of principal neurons in the hippocampus have a major role in gating synaptic plasticity at afferent synapses on proximal dendrites. Apical dendrites also receive powerful feed-forward inhibition mediated in large part by neurogliaform neurons. Here we show that theta-burst activity in afferents from the entorhinal cortex induces ‘Hebbian’ long-term potentiation at excitatory synapses recruiting these GABAergic cells. Such LTP increases disynaptic inhibition of principal neurons, thus shifting the excitation-inhibition balance in the temporoammonic pathway in favor of inhibition, with implications for computations and learning rules in proximal dendrites.

## Introduction

Regenerative potentials in the apical dendrites of pyramidal neurons have a profound effect on plateau potentials, spiking and synaptic plasticity in more proximal dendritic segments (Jarsky et al., 2005; Dudman et al., 2007; Takahashi and Magee, 2009; Basu et al., 2013; Han and Heinemann, 2013), and have been implicated in the formation of new CA1 place fields (Bittner et al., 2015). Computational studies have suggested that interactions between distal and proximal dendrites can support the multiplexing of information, and provide a plausible implementation of the backpropagation algorithm for supervised learning (Körding and König, 2001; Urbanczik and Senn, 2014; Guerguiev et al., 2017; Naud and Sprekeler, 2018; Sacramento et al., 2018; Richards and Lillicrap, 2019; Lillicrap et al., 2020; Payeur et al., 2021).

Importantly, the ability of excitatory afferents to distal dendrites to elicit regenerative events depends on the strength of disynaptic inhibition. In the hippocampus, the temporoammonic (TA) pathway from the entorhinal cortex (EC) powerfully recruits feedforward inhibition of apical dendrites in stratum lacunosum-moleculare (SLM). Indeed, bursts of TA activity can block both CA1 pyramidal cell spiking (Dvorak-Carbone and Schuman, 1999; Remondes and Schuman, 2002) and induction of long-term plasticity (LTP) (Levy et al., 1998; Remondes and Schuman, 2002) in response to Schaffer collateral stimulation, effects that depend on GABAergic transmission. Given the importance of feedforward inhibition of distal dendrites, it is pertinent to ask whether excitatory synapses that recruit such interneurons are plastic, and if so, whether they exhibit ‘Hebbian’ NMDA-receptor dependent LTP as reported in stratum radiatum interneurons (Lamsa et al., 2005, 2007a), rather than other forms of plasticity reported in parvalbumin-positive interneurons in stratum oriens (Perez et al., 2001; Lamsa et al., 2007b; Oren et al., 2009). Apical dendrites in CA1 also receive a direct innervation from the higher order thalamic nucleus reuniens (NRe) (Dolleman-van der Weel et al., 2019), which also recruits feedforward inhibition (Dolleman-Van der Weel et al., 1997; Dolleman-Van der Weel and Witter, 2000). A further question, therefore, is whether plasticity rules differ between synapses made by cortical and subcortical afferents.

While a heterogenous population of interneurons resides within SLM, or on the border between SLM and stratum radiatum (Vida et al., 1998; Klausberger, 2009; Capogna, 2011), the neurogliaform (NGF) subtype is the most abundant (Bezaire and Soltesz, 2013), and likely has a dominant role in SLM feedforward inhibition. NGF cells are innervated both by TA afferents and fibers from NRe (Chittajallu et al., 2017). They signal via volume transmission (Oláh et al., 2009), evoking exceptionally long-lasting inhibitory responses mediated by both GABA_A_ and GABA_B_ receptors (Price et al., 2008), and closely mimicking the effect of electrical stimulation of SLM (Williams and Lacaille, 1992). These postsynaptic response kinetics make NGF cells particularly well-suited to regulating dendritic nonlinearities and burst firing in pyramidal cells (Schulz et al., 2018). Interest in the properties and roles of NGF cells (Overstreet-Wadiche and McBain, 2015) has accelerated with the creation of a mouse line selective for neocortical layer 1 NGF cells, Ndnf-Cre (Tasic et al., 2016), although this has not, to date, been characterized extensively in the hippocampus.

Here, we first identify and characterize LTP mechanisms in SLM feedforward interneurons, and show, using an optogenetic strategy, that LTP and short-term facilitation can be induced at excitatory inputs from the EC but not at those from the thalamus. We confirm that the Ndnf-Cre mouse line can be used to target hippocampal NGF cells, and use it to demonstrate that this inhibitory cell subtype exhibits LTP in response to a physiologically relevant pattern of afferent activity. We further show that plasticity of feed-forward inhibition from the EC is not matched by parallel potentiation at EC-CA1 distal dendrite pyramidal cell synapses, and therefore that it alters the excitation-inhibition balance in the TA pathway.

## Materials and Methods

### Animals

Hippocampal slices were prepared from male and female wild-type mice (C57BL/6, Harlan) that were postnatal day 14 – 25 for electrically induced LTP experiments (Figs. 1 & 2), or 2 – 5 months old for optogenetic experiments (Fig. 3). Heterozygous Ndnf-Cre breeding pairs were obtained from The Jackson Laboratory (B6.Cg-Ndnf^tm1.1(folA/cre)Hze^/J; Stock No: 028536), and bred on a C57BL/6 background. Male and female 3 – 5 month old Ndnf-Cre mice were used for NGF cell LTP experiments (Fig. 4). All mice were housed under a non-reversed 12 h:12 h light-dark cycle and had access to food and water *ad libitum.* All procedures were performed in accordance with the UK Home Office Animals (Scientific Procedures) Act 1986.

**Fig. 1.**
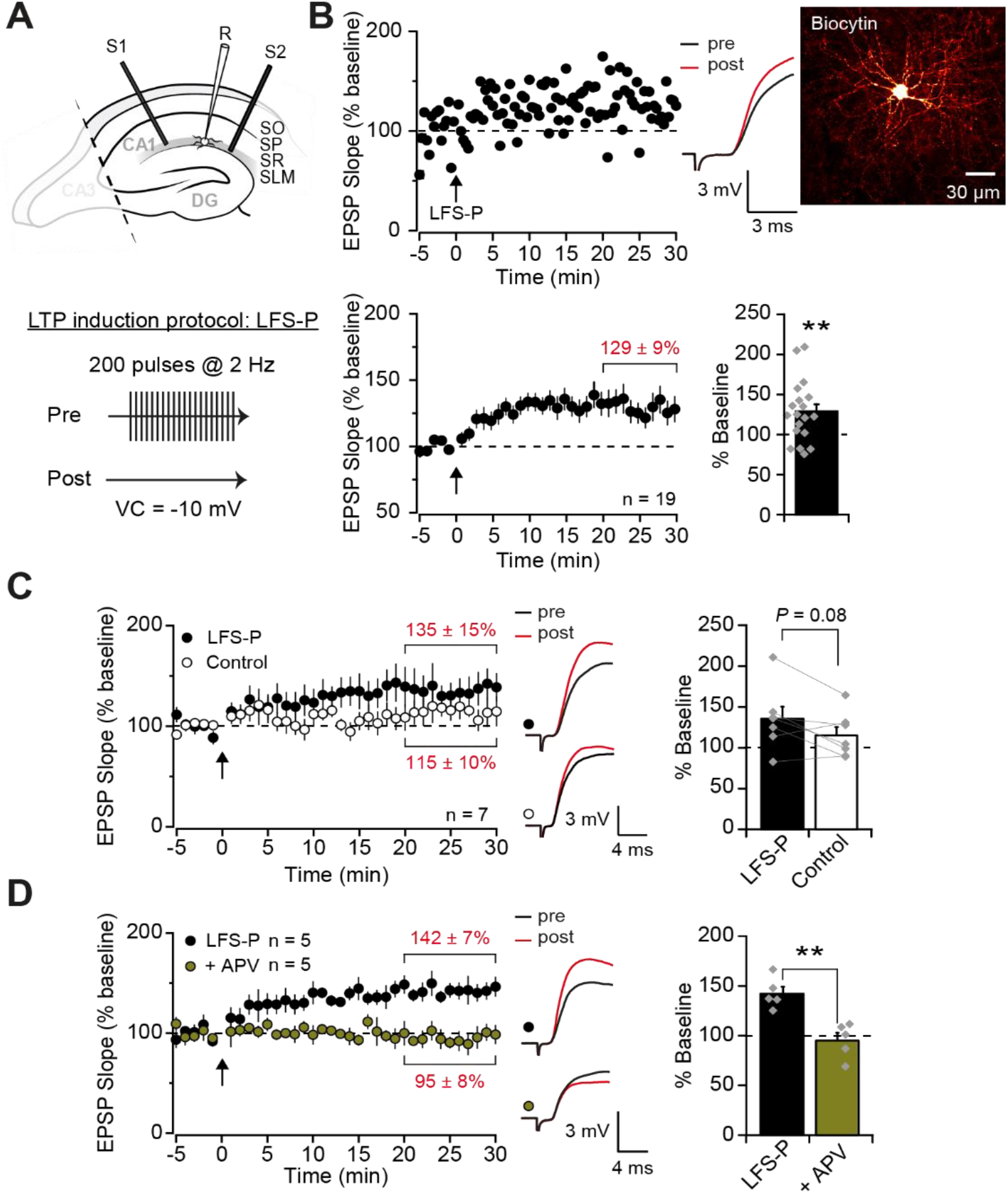
NMDA receptor-dependent LTP in SLM interneurons. (**A**) Top: experimental setup showing placement of one recording (R) and two stimulating (S1, S2) electrodes in SLM of a hippocampal slice with CA3 removed. Bottom: schematic of the low-frequency stimulation-pairing (LFS-P) LTP induction protocol. VC = voltage clamp. (**B**) Representative experiment and traces (top left) in a biocytin filled NGF-like cell (top right) and pooled dataset (bottom) showing LTP in SLM interneurons (quantified bottom right). Arrows indicate LTP induction protocol. (**C**) LFS-P-induced LTP was largely pathway-specific, quantified on right. (**D**) Application of APV (100 μM) blocked the induction of LTP by LFS-P, quantified on right. ** = *P* < 0.01. Error bars show standard error of the mean (SEM).

**Fig. 2.**
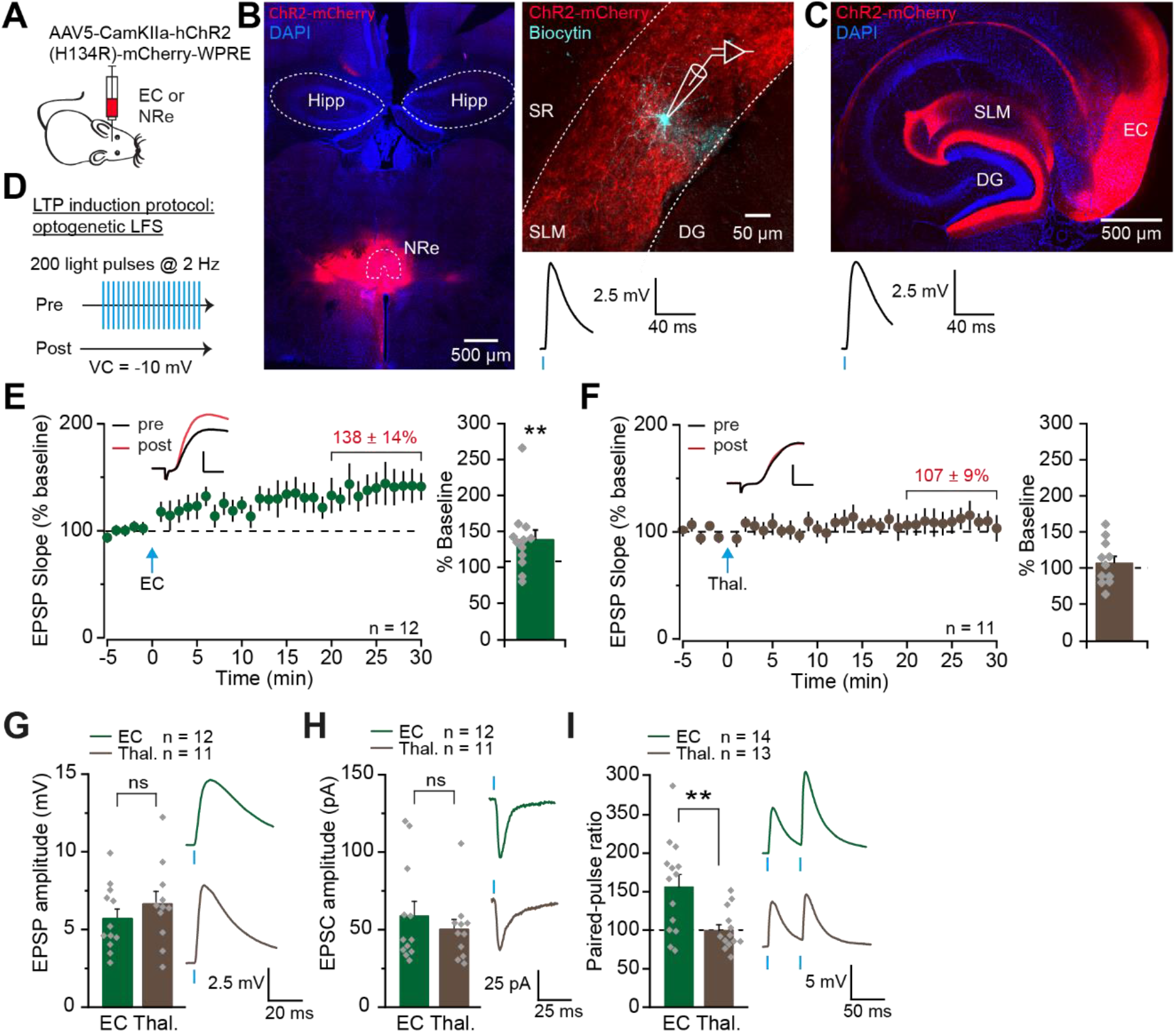
LTP induction by stimulation of entorhinal cortex but not thalamic inputs to SLM interneurons. (**A**) Schematic showing viral strategy used to target the nucleus reuniens (NRe) of the thalamus or the entorhinal cortex (EC). (**B**) Confocal images of a coronal brain section (70 μm) showing viral expression (red) at the injection site in the thalamus including the NRe, and DAPI staining (blue; left), and a sagittal hippocampal slice (300 μm) showing a biocytin-filled SLM interneuron (cyan) surrounded by transduced thalamic fibers (red; right). Below: example light-induced EPSP recorded from the same cell (average 15 traces). Hipp. = hippocampus, SR = stratum radiatum, DG = dentate gyrus. (**C**) Confocal image of a horizontal brain section (70 μm) showing viral expression (red) at the injection site in the EC, as well as in perforant path fibers in the SLM region of hippocampal CA1 and molecular layer of the dentate gyrus (DG), and DAPI staining (blue). Below: example light-induced EPSP recorded from an SLM interneuron in a mouse injected in the EC (average 15 traces). (**D**) Schematic of the optogenetic low-frequency stimulation-pairing (LFS-P) LTP induction protocol. (**E**) Optogenetic LFS-P of EC fibers induced LTP (scale: 3 mV, 4 ms), quantified on right. (**F**) The same optogenetic stimulation protocol applied to thalamic fibers did not induce LTP (scale: 3 mV, 4 ms), quantified on right. (**G**) Optogenetic stimulation of EC or thalamic fibers evoked EPSPs of similar amplitude, recorded during the baseline period of LTP experiments. (average of 15 traces). (**H**) As (A) but for EPSCs recorded during the optogenetic LFS-P LTP induction protocol (average 200 traces). (**I**) Paired-pulse facilitation (PPF) was induced by optogenetic paired-pulse stimulation of EC but not thalamic fibers (average 5 traces). ** = *P* < 0.01. Error bars show SEM.

**Fig. 3.**
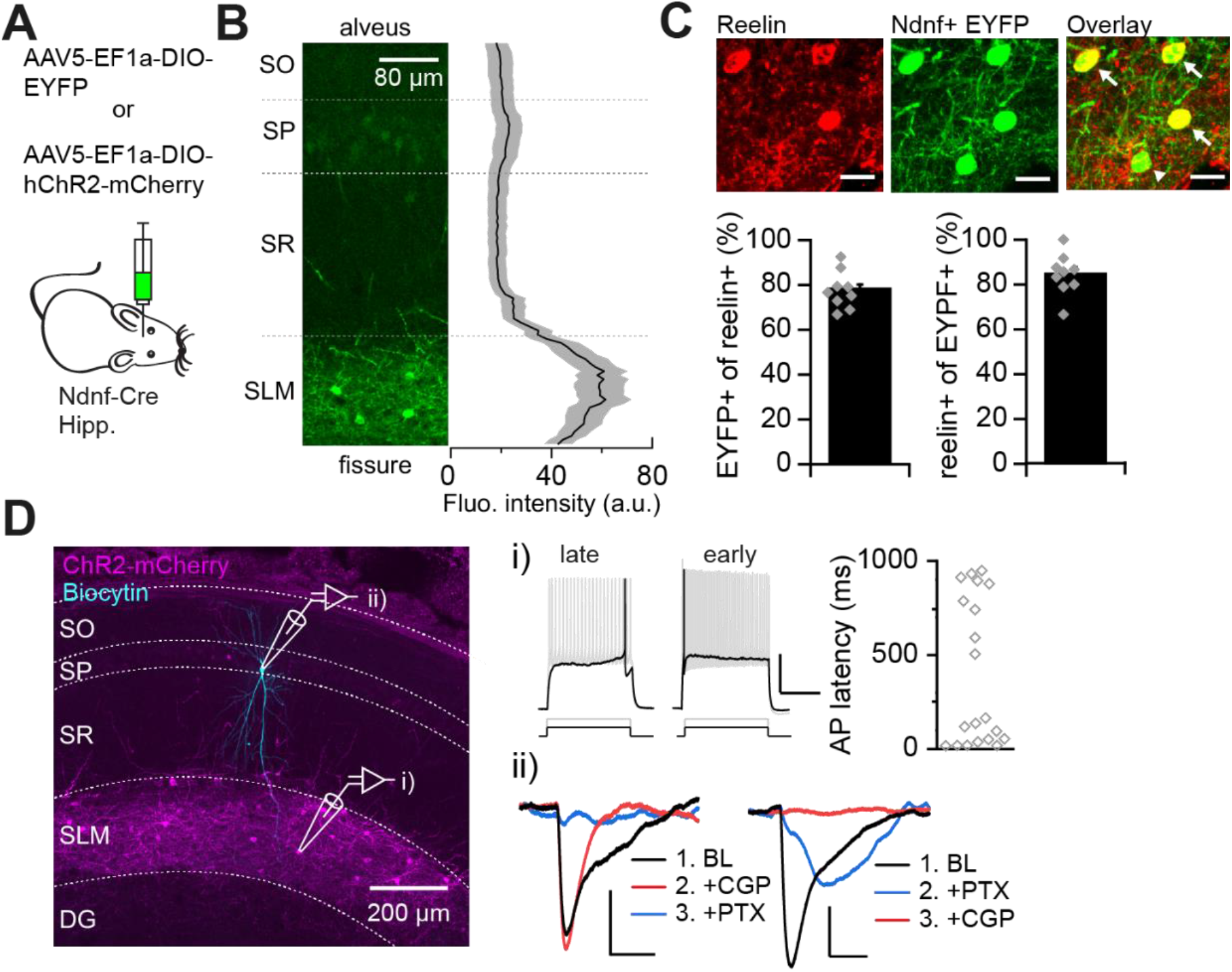
Hippocampal NDNF+ cells are NGF cells. (**A**) Schematic showing viral strategy used to express EYFP or ChR2-mCherry in hippocampal NDNF+ cells. Hipp. = hippocampus. (**B**) Left: confocal image of a section of CA1 used to measure NDNF+-EYFP fluorescence intensity across hippocampal layers. Right: mean (black) ± SEM (grey) fluorescence intensity across hippocampal layers (n = 6 slices, 4 mice), normalized to % distance from the alveus to the hippocampal fissure. SO = stratum oriens, SP = stratum pyramidale, SR = stratum radiatum. (**C**) Closeup of the SLM showing reelin immunostaining (top left), EYFP+ NDNF cells (top centre) and overlay (top right; scale: 20 μm). White arrows indicate reelin-positive NDNF cells and arrow head shows one reelin-negative NDNF cell. % overlap was quantified in 9 slices from 3 mice (bottom). (**D**) confocal image of a sagittal hippocampal slice (300 μm) showing selective expression in SLM neurons (magenta), and a biocytin-filled pyramidal cell (cyan) in stratum pyramidale. DG = dentate gyrus. (i) mCherry-tagged NDNF+ cells showed late- or early-spiking properties in response to a near-rheobase current step (black; quantified on right, n = 20), and sustained spiking in response to a double-rheobase current step (gray; scale: 25 mV, 500 ms). (ii) Optogenetic stimulation of NDNF+ cells induced long-lasting inhibitory postsynaptic potentials (IPSPs) in pyramidal cells at baseline (BL) (black) that were abolished by blockers of GABA_A_ (blue; PTX: picrotoxin, 100 μM) and GABA_B_ (red; CGP: CGP 55845, 1 μM) receptors (scales: 1 mV, 125 ms).

**Fig. 4.**
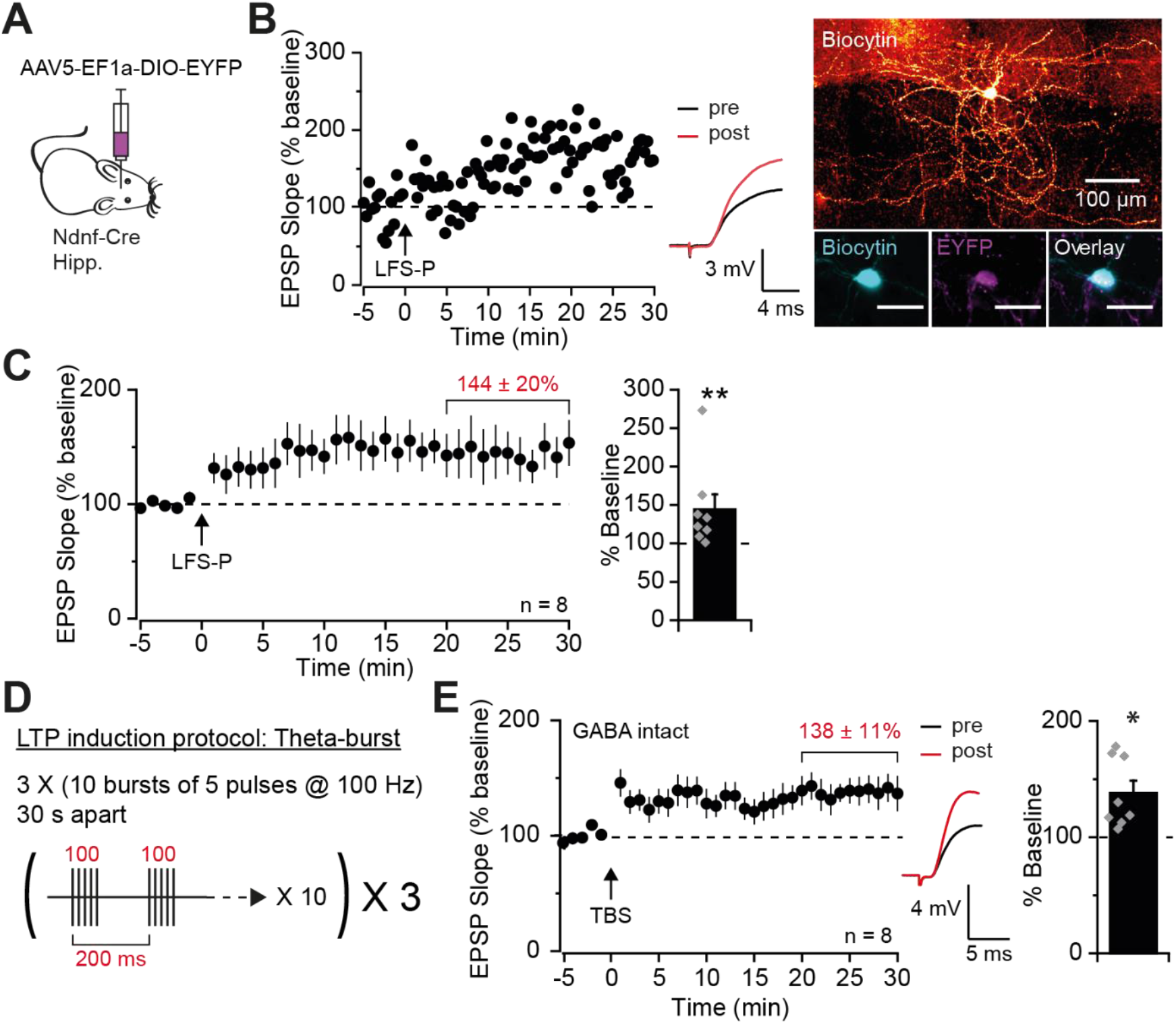
Low-frequency and theta-burst stimulation induced LTP in hippocampal NGF cells. (**A**) Schematic showing viral strategy used to express EYFP in hippocampal NDNF+ cells. Hipp. = hippocampus. (**B**) Representative experiment and traces of LFS-P-induced LTP (left) in a biocytin-filled EYFP-tagged NDNF+ cell (right; bottom right scale bars: 30 μm). (**C**) Pooled dataset showing LFS-P-induced LTP in NDNF+ NGF cells, quantified on right. (**D**) Schematic of the theta-burst stimulation (TBS) LTP induction protocol. (**E**) Pooled dataset showing TBS-induced LTP with GABAergic transmission intact in NDNF+ NGF cells, quantified on right. *\*P* < 0.05, \*\**P* < 0.01. Error bars show SEM.

### Surgery for viral injections

Mice (minimum age 6 weeks) were anesthetized using isoflurane and positioned in a stereotaxic frame, and on a heated pad to maintain body temperature. Either bilateral holes were made in the skull for hippocampal and EC injections, or a single hole was made in the midline for injections into the NRe. The coordinates used for each location were: EC, −4.3 mm caudal and +/-3.0 mm lateral of Bregma, and −3.2 and −3.0 mm deep from the pia with the syringe angled at 12 −14°; NRe, −0.7 mm caudal, on the midline, and −4.4, −4.2 and −4.0 mm deep; dorsal CA1 (SLM): −2.0 mm caudal and +/-1.25 mm lateral, −1.65 and −1.45 mm deep. The viral suspension (200 nl) was injected at each site (100 nl/min) through a 33 gauge needle using a Hamilton syringe, and the needle was left for 5 min post-injection before being withdrawn. The viruses used were AAV5-CaMKIIα-hchR2(H134R)-mCherry-WPRE, AAV5-EF1α-DIO-EYFP and AAV5-EF1a-DIO-hChR2-mCherry (all titres > 10^12^ viral genomes/ml) and were purchased from UNC Vector Core. Animals were monitored for ~5 days following the procedure, and acute experiments were performed a minimum of 3 weeks post-surgery to allow for viral expression.

### Slice preparation

Young mice (P14 - 25) were decapitated under isoflurane anesthesia and the brains were rapidly removed and placed in ice-cold, sucrose-rich slicing solution composed of (in mM): sucrose (75), NaCl (87), KCl (2.5), NaH_2_PO_4_ (1.25), NaHCO_3_ (25), Glucose (25), CaCl_2_ (0.5), MgCl_2_ (7), and oxygenated with 95% O_2_ and 5% CO_2_. Older mice (≥ 2 months) were given a lethal dose of sodium pentobarbital, and transcardially perfused with ice-cold oxygenated sucrose solution, before removal of the brain. 300 μm-thick sagittal slices were cut, except for a subset of brains that were cut in a horizontal orientation; results from experiments in sagittal and horizontal slices did not differ and were pooled for analysis. Slices were cut in ice-cold sucrose solution using a vibrating microtome (Leica VT1200 S), and were left to recover for 15 min at 32°C before being transferred to a standard artificial CSF (aCSF) solution containing (in mM): NaCl (119), KCl (2.5), MgSO_4_ (1.3), NaH_2_PO_4_ (1.25), NaHCO_3_ (25), Glucose (11), CaCl_2_ (2.5), and oxygenated with 95% O_2_ and 5% CO_2_. Slices were allowed to recover for at least 1 h at room temperature before being transferred to a recording chamber for experiments. The CA3 region was removed from slices in experiments where GABA receptors were blocked to prevent recurrent activity.

### Electrophysiology

#### Recording

Slices were held in place in a submerged recording chamber continuously perfused with oxygenated aCSF at a rate of 2-3 ml/min and maintained at 30-32°C. Picrotoxin (100 μM) and CGP55845 (1 μM) were routinely added to block GABA_A_ and GABA_B_ receptors, respectively, unless stated otherwise. Slices were visualized using an upright microscope (BX51WI, Olympus) and differential interference contrast (DIC) optics were used to identify cells located in the SLM layer of CA1. In wild-type mice, cells with small (~10 μm), round somata were selected for experiments, in line with known NGF cell morphology (Price et al., 2005). In Ndnf-Cre mice, EYFP- or mCherry-tagged NDNF+ cells were identified using blue (470 nm) or orange (590 nm) LED lights, respectively (ThorLabs) delivered through the microscope objective (20X; Olympus). Whole cell patch clamp recordings were performed using borosilicate glass micropipettes (2 – 4 MΩ) containing, for current clamp experiments, a K^+^-based internal solution containing (in mM): K-gluconate (125), KCl (5), KOH-HEPES (10), MgCl_2_ (1), Na-phosphocreatine (10), EGTA (0.2), Mg-ATP (4), Na_3_-GTP (0.4), and for voltage clamp experiments a cesium based solution containing (in mM): Cs-gluconate (125), NaCl (8), CsOH-HEPES (10), EGTA (0.2), Mg-ATP (4), Na-phosphocreatine (10), Na_3_-GTP (0.3), TEA-Cl (5). Solutions were adjusted to pH 7.2-7.5 and 290-295 mOsm, and biocytin (0.4%) was added for post-hoc morphological analysis in a subset of experiments. Recordings were made using a Multiclamp 700B amplifier (Molecular Devices), filtered at 10 kHz and digitized at 20 kHz, and experiments were run using custom LabVIEW (National Instruments) virtual instruments. Recordings were not corrected for liquid junction potential, and cells were rejected if they had an access resistance >25 MΩ, or access or input resistance that changed by more than 20% over the course of an experiment. Recordings for the low frequency stimulation pairing (LFS-P), some of the theta burst stimulation (TBS) (Fig. 4 & Extended Data Fig. 5-1), and the spike-timing dependent plasticity (STDP) experiments were made in current clamp mode at −70 mV, except during the LFS-P induction protocol when cells were temporarily switched to voltage clamp and depolarized to −10 mV. Disynaptic IPSCs and monosynaptic EPSCs in pyramidal cells were recorded in voltage clamp at +10 mV and −70 mV respectively. For LTP of monosynaptic EPSCs, cells were temporarily switched to current clamp for the duration of the TBS protocol.

#### Stimulation

Concentric bipolar stimulating electrodes (FHC) were positioned in the SLM and stimulation was delivered at 0.05 Hz via constant current stimulators (Digitimer). In experiments where 2 pathways were tested, stimulating electrodes were positioned on either side of the recorded cell, at opposite ends of SLM, and pathway independence was verified by comparing responses to paired-pulse stimulation within and across pathways: paired-pulses with an inter-pulse interval of 50 ms were applied to each pathway and typically revealed paired-pulse facilitation (PPF) ((EPSP1 slope/EPSP2 slope) x 100; average ± SEM: 176 ± 5%; n = 58). Paired-pulses were then applied across pathways, and pathways were considered independent if responses were not facilitated following stimulation of the other pathway. It should be noted that spikes were often triggered by larger EPSPs, thus preventing the measurement of PPF and pathway overlap in some cells; however, all pathways that could be tested were found to be independent (n = 29 pairs), suggesting that our stimulating electrode placement generally activated different sets of fibers. During experiments, stimulation was delivered to each pathway alternately, or to a single pathway, at a frequency of 0.05 Hz. Stimulation intensity for all electrically-induced LTP experiments was set to elicit sub-maximal EPSPs measuring ~5 mV in amplitude. For optogenetic stimulation of thalamic or EC fibers in LFS-P experiments, 1 ms-long pulses of blue light (470 nm) were delivered through the microscope objective to elicit EPSPs of amplitude ≥ 2.5 mV. Optogenetically elicited disynaptic IPSCs measured between 100 and 400 pA. Optogenetic stimulation for these experiments was delivered at 0.03 Hz to prevent use-dependent depression of NGF cell responses over time (Price et al., 2005).

Electrical or optogenetic LTP induction protocols were as follows: LFS-P: 200 pulses delivered at 2 Hz, while holding the cell at −10 mV in voltage clamp; STDP: 30 pulses delivered at 0.2 Hz, paired with 5 ms-long depolarizing current steps to induce 1 AP in the postsynaptic cell ~10 ms after presynaptic stimulation (cells were maintained at −65 mV in current clamp between current steps throughout this protocol to ensure sufficient depolarization for AP firing); TBS: 10 bursts of 5 pulses at 100 Hz delivered at theta frequency (5 Hz), and repeated 3 times, 30 s apart, while the postsynaptic membrane potential was allowed to float.

Electrophysiological properties of cells were assessed by injection of 1 s-long current steps, ranging from −200 pA to +300 pA in 50 pA increments. Rheobase current was also identified (minimal current injection required to evoke 1 AP), and a 1 s-long current step at twice rheobase delivered in order to assess firing frequency and adaptation; the latter was measured as the ratio between the firing frequency observed during the last 200 ms and that seen during the first 200 ms of the current step. Input resistance was measured and averaged from the −50 and +50 pA steps, and membrane time constant (τ) was calculated by fitting a single exponential function to the first 150 ms of the −50 pA step. Sag was calculated in two ways from the −200 pA current step: 1) as an absolute measure (mV) of the negative deflection seen at the start of the step, taken as the difference between the peak negative value and the steady state voltage measured at the end of the step (last 300 ms), and 2) this absolute value expressed as a percentage of the total negative deflection evoked by the −200 pA current injection, to take into account differences in input resistance across cells. AP and afterhyperpolarization (AHP) properties were calculated from the single AP evoked at rheobase current.

### Cell fixation and histochemistry

Cells were filled with biocytin during whole-cell recordings and slices were placed in 4% paraformaldehyde (PFA) overnight at 4°C. Fixed slices were rinsed 3 times in phosphate buffered saline (PBS), and subsequently stored in PBS containing 0.02% sodium azide. Biocytin was revealed by incubating slices in PBS with 0.3% Triton-X and streptavidin-conjugated Alexa Fluor (488 or 594, 1:1000; ThermoFisher Scientific, 21832 or S11227) for 3 h at room temperature. Some of the slices from Ndnf-Cre mice that had been injected with AAV5-EF1α-DIO-EYFP were also stained for GFP to enhance the EYFP-tag and confirm the identity of recorded cells. For these stainings, slices were first incubated in blocking solution consisting of 0.5% Triton-X, 0.5% bovine serum albumin (BSA) and 10% goat serum in PBS for 1 h at room temperature, before being transferred to PBS containing 0.5% Triton-X, 0.5% BSA and guinea pig anti-GFP antibody (1:1000; Synaptic Systems, 132 005) and left overnight at 4°C. After washing in PBS, slices were incubated in goat anti-guinea pig secondary antibody conjugated with Alexa Fluor 488 (1:500; ThermoFisher Scientific, A-11073) and streptavidin-conjugated Alexa Fluor 594 (1:1000; ThermoFisher Scientific, S1127). DAPI staining was then performed by incubating slices for 5 min in PBS with DAPI (1:5000).

For characterization of reelin expression, Ndnf-Cre mice injected with AAV5-EF1α-DIO-EYFP were transcardially perfused with 4% PFA and brains were removed and post-fixed in the same solution for at least 24 h. After rinsing with PBS, brains were sliced coronally (70 μM) and reelin staining was performed as described above, using mouse anti-reelin (1:1000; Abcam, ab78540) and goat anti-mouse Alexa Fluor 568 (1:500; Invitrogen A-11004). Expression was quantified in 3 hippocampal sections from 3 injected mice (n = 9): the SLM was manually selected as the region of interest (ROI) in each slice and reelin+ and NDNF+ EYFP cells were counted separately before overlaying images to quantify overlap.

### Experimental design and statistical analysis

For LTP experiments, *n* values represent individual cells from separate slices, obtained from at least 3 mice per experiment. All experiments involving pharmacological manipulations were interleaved with control experiments. Individual experiments were analyzed using custom code written in Python, and statistical analysis was carried out in Python and Origin (2018; OriginLab). For quantification of LTP, paired students *t*-tests were carried out on raw pre- (baseline) vs post-LTP (last 10 min) EPSP slope or amplitude measurements. EPSP slope was measured during a 2 ms window from the onset of the EPSP. Test and control pathways were compared using paired *t*-tests on normalized responses averaged from the last 10 min of experiments, and unpaired *t*-tests were used to compare pharmacological manipulations and optogenetic responses to thalamic versus EC stimulation. Shapiro-Wilk tests were used to assess normality, and paired and unpaired *t-*tests were replaced by Wilcoxon signed rank and Mann-Whitney U tests, respectively, if data were found to be non-normally distributed. Spearman rank correlation was used to assess correlations. Differences were considered significant when *P* < 0.05, and are indicated in the figures as *P* < 0.001 (***), *P* < 0.01 (**), *P* < 0.05 (*), or *P* = X when 0.05 < *P* < 0.1. All data are presented as mean ± SEM. Representative traces are an average of 15 traces taken during the 5 min baseline period or during the last 5 min of recordings, unless stated otherwise.

## Results

### LTP in SLM interneurons

We recorded from interneurons in CA1 SLM with small round somata, consistent with NGF cell morphology (Price et al., 2005), and stimulated afferent fibers in the same layer (Fig. 1*A*). We restricted attention to the initial slope of EPSPs and employed a low-frequency stimulation (LFS) protocol paired with postsynaptic depolarization (LFS-P; Fig. *1*A) in order to look for LTP at monosynaptic excitatory inputs (Maccaferri and McBain, 1996). Inhibitory transmission was blocked by inclusion of GABA_A_ and GABA_B_ receptor antagonists in the aCSF. LFS-P led to an increase in EPSP slope to 129 ± 9% of baseline, lasting for at least 30 minutes (Fig. 1*B*; n = 19; *t*(18) = 3.47, *p* = 0.003, paired *t*-test). EPSP slopes after LFS-P ranged from 77% to 210% of baseline (Fig. 1*B*). Neurogliaform cells in the neocortex tend to fire either early or late during threshold current injections (Jiang et al., 2015). The extent of potentiation did not correlate with the latency to the first action potential in the present dataset (Extended Data Fig. 1-1; n = 19; *r* = 0.19, *p* = 0.45, Spearman test), suggesting that differences in interneuron subtype did not underlie the observed variability in LTP magnitude. Dual pathway experiments revealed a larger increase in the paired pathway than in a control pathway that was not stimulated during the LFS-P protocol (Fig. 1*C*; test: 135 ± 15%, control: 115 ± 10%, n = 7), although this difference did not reach statistical significance (*t*(6) = 2.12, *p* = 0.08, paired *t*-test). This observation suggests that LTP may not be fully restricted to stimulated synapses, as might be expected in cells with aspiny dendrites (Cowan et al., 1998; Chen and Sabatini, 2012), such as SLM interneurons (Lacaille and Schwartzkroin, 1988) and NGF cells (Price et al., 2005).

Consistent with LTP in principal cells that depends on NMDA receptors, LFS-P-induced LTP in SLM interneurons was fully blocked by perfusion of (DL)-aminophosphonovalerate (APV, 100 μM; Fig. 1*D*; interleaved LFS-P: 142 ± 7%, n = 5; APV: 95 ± 8%, n = 5; *t*(8) = 4.49, *p* = 0.002, unpaired *t*-test). Importantly, in control experiments neither presynaptic LFS alone nor postsynaptic depolarization alone led to potentiation, further confirming its Hebbian nature (Extended Data Fig. 1-2*A*: LFS alone: 96 ± 15%, n = 6; *t*(5) = 0.23, *p* = 0.83, paired *t*-test; Extended Data Fig. 1-2*B*: depolarization alone: 111 ± 9%, n = 5; *t*(4) = −1.32, *p* = 0.26, paired *t*-test).

In contrast to LFS-P, a spike timing-dependent plasticity (STDP) protocol consisting of 30 presynaptic stimuli, each followed by a single postsynaptic spike (Extended Data Fig. 1-3*A*) induced a potentiation that resisted NMDA receptor blockade (Extended Data Fig. 1-3*B*; STDP: 136 ± 8%, n = 15; *t*(14) = −4.55, *p* <0.001, paired *t*-test; APV: 135 ± 11%, n = 6; *t*(5) = 3.55, *p* = 0.02, paired *t*-test). Instead, this was blocked by the voltage-gated Ca^2+^ channel blockers nimodipine (10 μM) and Ni^2+^ (100 μM) (Extended Data Fig. 1-3*C*; 109 ± 13%, n = 5; *t*(4) = −0.39, *p* = 0.71, paired *t*-test), indicating that NMDA receptor-independent Ca^2+^ influx, likely triggered by back-propagating action potentials interacting with EPSPs, can also be sufficient for LTP induction in these cells, as previously reported in other hippocampal interneurons (Galván et al., 2008; Nicholson and Kullmann, 2017).

### LTP induction by stimulation of EC but not thalamic inputs to SLM interneurons

The SLM region of CA1 receives excitatory projections from both layer III of the EC (Witter et al., 1988) and the NRe of the thalamus (Wouterlood et al., 1990). Electrical stimulation in SLM however does not distinguish between afferents from these regions, which are likely to have distinct functional roles in modulating hippocampal activity. In order to activate either pathway selectively, we injected an adeno-associated virus (AAV) encoding channelrhodopsin-2 (ChR2) under the CaMKII promoter into either the NRe or the EC (Fig. 2*A-C*), and recorded responses in SLM interneurons to optogenetic stimulation of ChR2-expressing axons (Fig. 2*B* and *C*). Importantly, viral injections in either location led to expression of ChR2 within the hippocampus that was restricted to SLM (Fig. 2*B* and *C*), and, in the case of EC injections, the molecular layer of the dentate gyrus (Fig. 2*C*), as shown previously (Chittajallu et al., 2017). Light stimulation reliably evoked EPSPs in SLM interneurons following expression of ChR2 in either set of fibers (Fig. 2*B* and *C*). However, optogenetic responses evoked by stimulation of thalamic (but not EC) fibers were not stable and instead gradually increased with time (Extended Data Fig. 2-1*A*: thalamus: 136 ± 19% after 30 mins recording, n = 3; Extended Data Fig. 2-1*B*: EC: 97 ± 5% after 30 mins recording, n = 13), precluding the ability to measure LTP of these responses. The cause of this ‘run-up’ is unclear but could be due to progressive Ca^2+^ loading of presynaptic terminals following ChR2-induced action potentials (Zhang and Oertner, 2007). For this reason, light stimulation was used to deliver a conditioning stimulus selectively to either EC or thalamic terminals, whilst electrical stimulation was used to monitor EPSPs at synapses made by both sets of afferents. Pairing optogenetic stimulation of EC inputs with depolarization, in a LFS-P protocol identical to that used in Fig. 1 but with electrical stimulation interrupted (Fig. 2*D*), led to an increase in EPSP slope to 138 ± 14% of baseline (Fig. 2*E*; n = 12; *t*(11) = 3.85, *p* = 0.003, paired *t*-test). In contrast, the same pairing applied to thalamic afferents had no observable effect, with the EPSP slope remaining at 107 ± 9% of baseline (Fig. 2*F*; n = 11). Notably, light-evoked responses were of similar amplitude whether stimulating EC or thalamic terminals, both during baseline recordings of EPSPs in current clamp (Fig. 2*G*; EC: 5.7 ± 0.6 mV, n = 12; thalamus: 6.6 ± 0.8 mV, n = 11; *t*(21) = 0.93, *p* = 0.36, unpaired *t*-test), and during the LTP induction protocol in voltage clamp (Fig. 2*H*; EC: 59.0 ± 9.3 pA, n = 12; thalamus: 50.2 ± 6.4 pA, n = 11; *U* = 74, *p* = 0.64, Mann-Whitney U test), suggesting that the different outcomes could not be explained by the degree of SLM interneuron depolarization. Interestingly, however, optogenetically evoked responses to stimulation of EC fibers, but not thalamic fibers, exhibited paired-pulse facilitation (PPF) (Fig. 2*I*; EC: 156 ± 16%, n = 14; thalamus: 100 ± 7%, n = 13; *t*(25) = 3.10, *p* = 0.005, unpaired *t*-test), consistent with a lower release probability in the TA pathway. The differences between these pathways potentially contribute to the variability in LTP magnitude seen with electrical induction (Fig. 1) if the ratio of EC and NRe afferents varies depending on the position of the stimulating electrode.

### Hippocampal NDNF+ cells correspond to NGF cells

NGF cells constitute ~10% of the total inhibitory neuron population in hippocampal area CA1, and the overwhelming majority are located in SLM (Bezaire and Soltesz, 2013). The neurons recorded from in the present study had a somatic shape typical of NGF cells. Indeed, biocytin filling and post-hoc morphological reconstruction confirmed that at least some of the cells in which LTP was studied were indeed NGF interneurons (Fig. 1*B*). However, their fine dendritic and axonal arborization renders morphological recovery particularly challenging (Tremblay et al., 2016), preventing routine identification. In order to directly address whether hippocampal NGF cells express LTP, we used Neuron-Derived Neurotrophic Factor (Ndnf)-Cre mice, a mouse line shown to specifically target NGF cells in cortical layer I ((Tasic et al., 2016), see also (Abs et al., 2018; Schuman et al., 2019)). As NDNF has principally been identified as an NGF cell-specific marker in the neocortex, we first sought to characterize hippocampal expression of NDNF+ cells by injecting the dorsal hippocampus of Ndnf-Cre mice with an AAV encoding either Cre-dependent enhanced yellow fluorescent protein (EYFP) or Cre-dependent ChR2 tagged with mCherry (Fig. 3*A*). NDNF+ cells were abundant in the hippocampus, and were found almost exclusively in SLM (Fig. 3*B,D*; fluorescence intensity analyzed in 6 slices from 4 mice), consistent with the known location of NGF cells. Weak expression was occasionally observed in both CA1 and dentate gyrus cell body layers, but never in hippocampal strata where other interneuron subtypes are located, such as radiatum and oriens; this suggests a possible minor leak in excitatory cells, but likely selective targeting of NGF cells amongst other hippocampal interneuron subtypes. Indeed, immunostaining for reelin, a molecular marker selectively expressed by hippocampal NGF cells (Fuentealba et al., 2010), confirmed that the overwhelming majority of NDNF+ cells in the SLM were reelin-positive (Fig. 3*C*; 84.5 ± 3%, n = 9 slices from 3 mice). Furthermore, threshold current injections in NDNF+cells revealed two populations with either early- or late-spiking properties (Fig. 3*Di*, n = 20), matching the firing pattern described in both cortical NDNF+ (Tasic et al., 2016) and NGF (Jiang et al., 2015) cells; other electrophysiological properties were also within the ranges described for both hippocampal (Tricoire et al., 2010) and cortical NGF (Karagiannis et al., 2009; Jiang et al., 2015) cells, as well as NDNF+ cells ((Tasic et al., 2016) Extended Data Table 3-1). Finally, optogenetic activation of hippocampal NDNF+ cells produced long-lasting inhibitory responses in pyramidal neurons comprising a GABA_A-slow_ and a strong GABA_B_ component, evident both in the biphasic nature of the response and in the pharmacological dissection of its components (Fig. 3*Dii* and Extended Data Table 3-2), both of which are notable hallmarks of NGF cell signaling (Tamas, 2003; Price et al., 2005; Capogna and Pearce, 2011). Together, these properties strongly support the use of Ndnf-Cre mice for the selective targeting of hippocampal NGF cells.

### Hippocampal NGF cells exhibit LTP

Having confirmed the identity of hippocampal NDNF+ cells, we next sought to verify that LTP could be induced in NGF cells by recording from fluorescently tagged NDNF+ cells in SLM and applying the LFS-P protocol with electrical stimulation. This induced a clear increase in EPSP size to 144 ± 20% of baseline (Fig. 4*B* and *C*; n = 8; *Z* = 2.45, *p* = 0.008, Wilcoxon signed rank test), demonstrating that hippocampal NGF cells exhibit robust LTP. We then asked whether this form of plasticity occurs under more physiological conditions by excluding GABA receptor blockers from the aCSF to leave inhibitory transmission intact, and by applying a theta-burst stimulation (TBS) protocol to mimic activity patterns at EC-CA1 synapses (Buzsáki, 2002). Under these conditions, excitatory transmission onto NGF cells was significantly potentiated to 138 ± 11% of baseline (Fig. 4*E*; n = 8; *Z* = −2.45, *p* = 0.01, Wilcoxon signed rank test), indicating that excitatory inputs onto hippocampal NGF cells are likely to be potentiated by naturally occurring activity.

### SLM feedforward interneuron LTP alters E-I balance at EC-CA1 synapses

What effect does LTP at synapses on SLM interneuron have on the downstream hippocampal network? We first asked whether potentiation of excitatory transmission onto SLM interneurons translates into greater feedforward inhibition of CA1 pyramidal cells. The TBS protocol is not only physiologically relevant but also avoids the need for experimentally imposed depolarization of postsynaptic cells via a patch clamp pipette, and should therefore allow LTP to be induced in a population of interneurons mediating disynaptic inhibition. We held CA1 pyramidal cells in voltage clamp mode at +10 mV and recorded inhibitory postsynaptic currents (IPSCs) evoked by electrical stimulation in SLM (Fig. 5*A*). To assess whether IPSCs were disynaptic or contaminated by direct activation of interneurons, and therefore monosynaptic, we blocked AMPA receptors with NBQX (10 μM). This however failed to suppress IPSCs generated by electrical stimulation, even when the stimulating electrode was placed as far from the recorded pyramidal cell as possible (Fig. 5*B*; 103 ± 8 %, n = 4; *t*(3) = −0.23, *p* = 0.83, paired *t*-test). This finding is in line with a previous study showing that electrical stimulation of the SLM as far as 900 μm from the recorded pyramidal cell directly recruits GABAergic neurons mediating monosynaptic inhibition (Milstein et al., 2015). We therefore switched to an optogenetic strategy to elicit disynaptic inhibition of CA1 pyramidal neurons, and confirmed that IPSCs evoked by light pulses following ChR2 expression in the EC were abolished by NBQX (Fig. 5*B*; 10 ± 8 %, n = 4; *t*(3) = 3.13, *p* = 0.05, paired *t*-test). When optogenetic stimulation of EC fibers was used to both elicit disynaptic inhibition and deliver the TBS protocol, LTP was reliably induced, detected as a significant increase in feedforward inhibition onto CA1 pyramidal cells (Fig. 5*C, D*; 135 ± 8%, n = 10; *t*(9) = −4.10, *p* = 0.003, paired *t*-test). Finally, we asked whether this form of LTP serves to maintain the excitation-inhibition (E-I) balance of the EC input in the face of potentiation taking place in parallel at excitatory inputs onto CA1 pyramidal cells, as has been shown for LTP of feedforward inhibition in stratum radiatum (Lamsa et al., 2005). In order to record monosynaptic EPSCs (similarly elicited by optogenetic EC stimulation), we held the pyramidal cell at −70 mV in voltage clamp, but switched to current clamp during the LTP induction protocol to allow it to depolarize naturally in response to the optogenetic TBS. Under these conditions, designed to match those that induce LTP of feedforward inhibition, we saw no potentiation of EPSCs in CA1 pyramidal cells (Fig. 5*D*; 99 ± 4%, n = 10; *t*(9) = −0.02, *p* = 0.98, paired *t*-test). Importantly, TBS of the SLM in the presence of GABA blockers induced robust potentiation in CA1 pyramidal cells (Extended Data Fig. 5-1; 184 ± 7%, n = 7; *t*(6) = −6.33, *p* < 0.001, paired *t*-test), indicating that LTP can occur at this synapse but is tightly regulated by feedforward inhibition. LTP at synapses on SLM interneurons thus alters the E-I balance in the EC input to hippocampal CA1 in favor of inhibition.

**Fig. 5.**
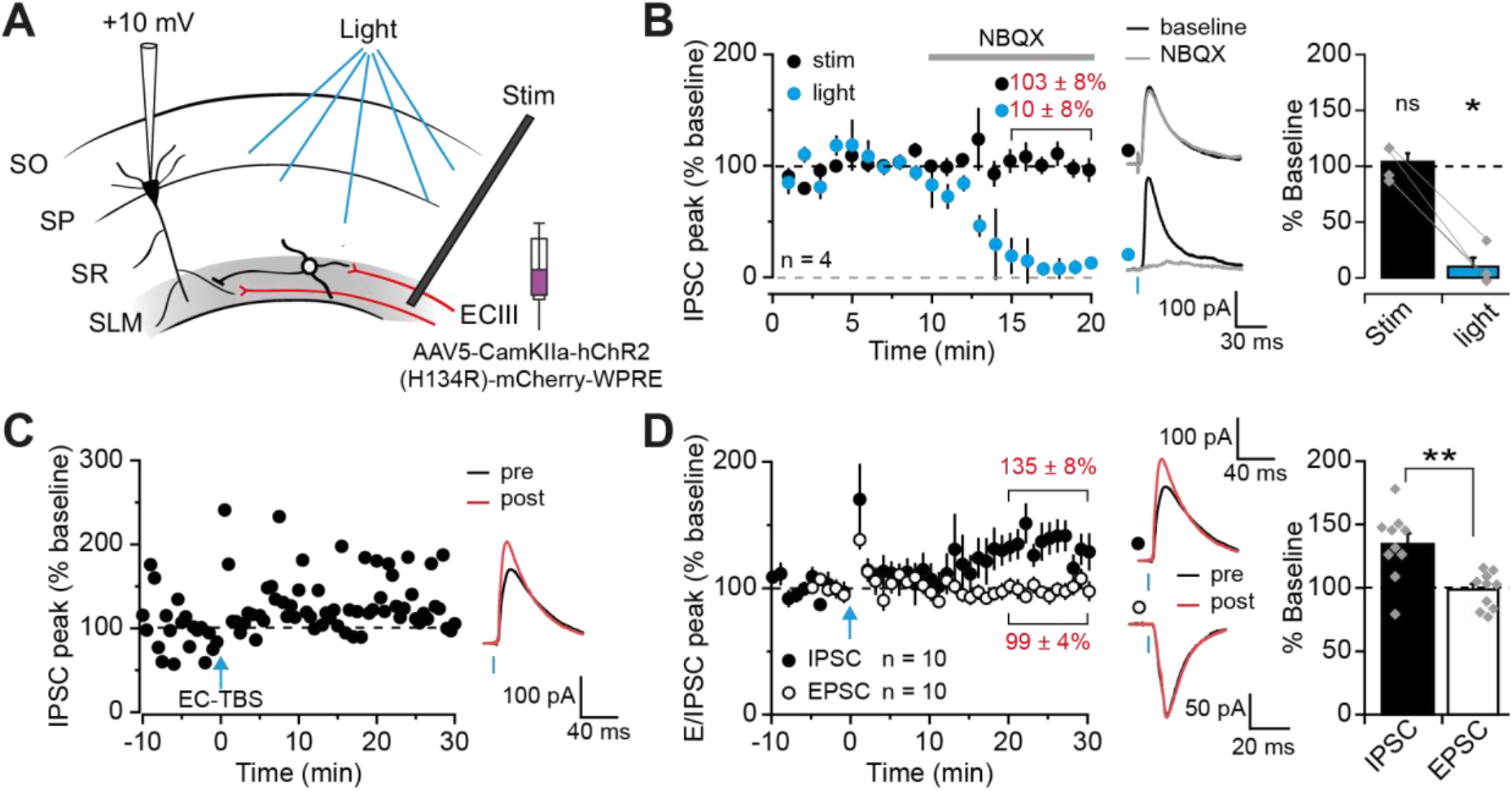
LTP of SLM interneurons alters E-I balance at EC-CA1 synapses. (**A**) Experimental setup for recording disynaptic IPSCs from CA1 pyramidal cells elicited by electrical stimulation of the SLM (stim) or optogenetic stimulation of EC fibers (light). SO = stratum oriens, SP = stratum pyramidale, SR = stratum radiatum. (**B**) Effect of NBQX (10 μM) on IPSCs elicited by alternate electrical and optogenetic stimulation. Reduction of IPSCs quantified on right. (**C**) Representative experiment and traces showing LTP of feedforward inhibition induced by optogenetic TBS of EC fibers (EC-TBS). (**D**) Pooled dataset showing EC-TBS induced LTP of disynaptic IPSCs (black), but not of monosynaptic EPSCs (white), with representative traces (IPSCs same as panel (C)). Quantified on right. \**P* < 0.05, \*\**P* < 0.01. Error bars show SEM.

## Discussion

The present study shows that interneurons located in SLM express Hebbian, NMDA receptor-dependent LTP, and that this can be induced by optogenetic stimulation of inputs originating in EC layer III but not, under the conditions tested here, by stimulation of afferents from the NRe of the thalamus. Importantly, we confirm that a mouse line previously developed to target cortical NGF cells, Ndnf-Cre (Tasic et al., 2016), also selectively labels hippocampal NGF cells, and show, using this novel tool, that this prominent interneuron subtype expresses LTP. Finally, using a theta-burst protocol resembling the natural activity of EC inputs to CA1, with inhibition intact, we show that plasticity is readily induced at EC-SLM interneuron synapses but not at monosynaptic connections from EC to pyramidal cell distal dendrites, and that this translates into a downstream shift in E-I balance in favor of inhibition.

The finding that CA1 interneurons located in SLM, of which NGF cells account for a large fraction, exhibit long-term plasticity has not, to our knowledge, previously been shown. Indeed, one earlier study reported no evidence of LTP, although it focused on interneurons located at the border of SLM and stratum radiatum with stimulation of excitatory fibers in both strata (Ouardouz and Lacaille, 1995). The present findings, however, complement work showing plasticity at mossy fiber inputs to SLM interneurons located in CA3 (Galván et al., 2008), as well as evidence of LTP in CA1 stratum radiatum Ivy cells (Szabo et al., 2012), a cell-type closely related to NGF cells (Tricoire et al., 2010; Armstrong et al., 2012), and of plasticity reported in cortical NDNF+ NGF cells (Abs et al., 2018). We found that LTP in SLM interneurons is typically NMDA receptor-dependent and at least partially pathway-specific. This is in line with previous reports of pathway-specific, NMDA receptor-dependent Hebbian LTP in aspiny hippocampal interneurons (Lamsa et al., 2005; Galván et al., 2015), and further challenges the view that spines are necessary for the dendritic compartmentalization of plasticity (Yuste et al., 2000; Nimchinsky et al., 2002). Interestingly, we found that an STDP protocol induced an NMDA receptor-independent form of plasticity in these cells, that was instead dependent on Ca^2+^ influx through VGCCs. This phenomenon is reminiscent of plasticity mechanisms seen in oriens-alveus interneurons, where NMDA receptor-independent LTP requires Ca^2+^-permeable AMPA receptors, T-type Ca^2+^ channels or nicotinic receptors (Nicholson and Kullmann, 2014, 2017, 2021). Together, these results argue that postsynaptic Ca^2+^ is necessary for LTP induction in SLM interneurons, but that the source of Ca^2+^ itself can vary.

Notably, selective optogenetic stimulation of EC fibers induced LTP in SLM interneurons whilst equivalent stimulation of thalamic fibers did not. This could be due to a number of pre- and/or post-synaptic differences between these two synapses, such as the specific receptors present, the probability of release at each synapse, and the resultant short-term plasticity mechanisms that may be at play during the induction protocol. In support of the latter hypothesis, optogenetic dissection of the two inputs revealed robust paired-pulse facilitation only at synapses made by afferents from the EC. Although short-term facilitation has previously been reported in response to NRe stimulation *in vivo* (Dolleman-Van der Weel et al., 1997, 2017; Bertram and Zhang, 1999; Gruart et al., 2015), these studies did not examine synapses on inhibitory cells. It is possible that NRe synapses on SLM interneurons exhibit a higher basal release probability than those onto pyramidal cells, thereby resulting in an absence of paired-pulse facilitation at the former. Indeed, previous studies have shown that neurotransmitter exocytosis probability can vary in a target cell-dependent manner, even across synapses formed by the same presynaptic axon (Koester and Johnston, 2005; Branco and Staras, 2009).

In the broader context of feedback connections and their computational function, the different short- and long-term plasticity rules revealed here at cortical and subcortical inputs from the EC and thalamus, respectively, are particularly interesting. Indeed, the plasticity displayed by EC inputs onto SLM interneurons corresponds to recent computational work proposing that hierarchically connected networks can coordinate learning via burst-dependent plasticity and multiplexing of top-down and bottom-up signals (Payeur et al., 2021). In this model, short-term facilitation is required for signal multiplexing, while LTP of inhibition may regulate pyramidal cell plasticity via control of burst probability. In contrast, the absence of either short- or long-term plasticity at NRe inputs, which are also thought to part of a higher-order cortico-thalamo-cortical circuit (Dolleman-van der Weel et al., 2019), suggests that alternative mechanisms may be involved in coordinating learning between these structures. Alternatively, neuromodulation, for instance mediated by local cholinergic inputs, may be required to reveal plasticity at NRe-SLM interneuron synapses, as has been suggested for stratum oriens interneurons (Nicholson and Kullmann, 2021). These open questions warrant further experimental and computational investigation.

Ndnf-Cre mice have, hitherto, primarily been used to label NGF cells located in cortical layer I (Tasic et al., 2016; Abs et al., 2018). Our results however indicate that this mouse line also enables selective targeting of NGF cells within the hippocampus, in line with previously reported expression of NDNF in this subcortical structure (Kuang et al., 2010). Thus, we show that hippocampal NDNF+ cells are located primarily within SLM, express the NGF cell-associated molecular marker reelin (Fuentealba et al., 2010), exhibit electrophysiological properties that are consistent with those previously described for both hippocampal (Tricoire et al., 2010) and cortical (Karagiannis et al., 2009; Jiang et al., 2015; Tasic et al., 2016) NGF cells, and induce slow, long-lasting GABA_A_ and GABA_B_ receptor-mediated postsynaptic responses typical of NGF interneuron signaling (Tamas, 2003; Price et al., 2005; Oláh et al., 2009; Capogna and Pearce, 2011). While these results strongly support selectivity for NGF cells amongst hippocampal interneurons, weaker expression was also occasionally observed in putative excitatory cells. Hippocampal NGF cells have previously been difficult to target or manipulate selectively. Indeed, whilst neuronal nitric oxide synthase (nNOS; (Taniguchi et al., 2011; Li et al., 2014; Bloss et al., 2016)) and neuropeptide Y (NPY; (Tricoire et al., 2010; Krook-Magnuson et al., 2011; Chittajallu et al., 2013; Li et al., 2017; Jackson et al., 2018)), amongst others, have been used as markers to aid in their identification, neither is fully selective for NGF cells (Overstreet-Wadiche and McBain, 2015; Pelkey et al., 2017). The Ndnf-Cre mouse line is thus the first genetic tool to achieve such selectivity, and will no doubt prove invaluable for investigations of hippocampal NGF cell function. Furthermore, a recent study showing that NDNF is a conserved marker for NGF cells in the human cortex (Poorthuis et al., 2018) implies that results obtained with this mouse line could be translated to humans.

The finding that TBS with intact GABAergic transmission, reminiscent of natural activity patterns and conditions found in the direct EC-CA1 pathway (Buzsáki, 2002), induces LTP in SLM interneurons and NGF cells, but not in pyramidal cells, is notable. On a mechanistic level, failure to elicit LTP of the excitatory TA pathway may be due to the distal location of EC inputs on pyramidal cell dendrites, which, when combined with powerful feedforward inhibition, limits the ability of the TA pathway to drive membrane depolarization in these cells. Importantly, the downstream effect is a net shift in E-I balance in favor of inhibition, in contrast to the effect of LTP in feedforward stratum radiatum interneurons reported previously (Lamsa et al., 2005). Indeed, LTP in these inhibitory cells was shown to counterbalance potentiation occurring in parallel at Schaffer collateral-CA1 pyramidal cell synapses, and thereby to maintain E-I balance and preserve the temporal fidelity of synaptic integration. Feedforward inhibition in the SLM is thought to impose a time window within which TA and Schaffer collateral inputs can interact nonlinearly (Capogna, 2011; Overstreet-Wadiche and McBain, 2015), and thus likely has a restrictive role on the generation of dendritic spikes (Jarsky et al., 2005), plateau potentials and plasticity (Remondes and Schuman, 2002; Bittner et al., 2015) in CA1 pyramidal cells. Interestingly, inhibition in layer I of the cortex, mediated largely by NGF cells (Jiang et al., 2015), has been proposed to have an analogous role in regulating dendritic nonlinearities generated by input coupling (Larkum et al., 1999), and activation of cortical NDNF+ NGF cells has been shown to inhibit the generation of dendritic spikes in layer V pyramidal neurons (Abs et al., 2018). It seems likely, then, that plasticity of excitatory inputs onto NGF cells, both within the hippocampus and in the neocortex (if indeed it occurs there), and the resultant shift in E-I balance, will lead to a tighter regulation of supralinear dendritic integration in pyramidal cells. Future experiments studying this phenomenon directly, and employing *in vivo* optogenetic strategies in Ndnf-Cre mice, will be instrumental in understanding the impact of potentiating TA feedforward inhibition on the wider hippocampal network and behavior.

## Acknowledgements

This work was supported by the Wellcome Trust, the Medical Research Council and Epilepsy Research UK (D.M.K).

## Extended data

**Extended Data Fig. 1-1.**
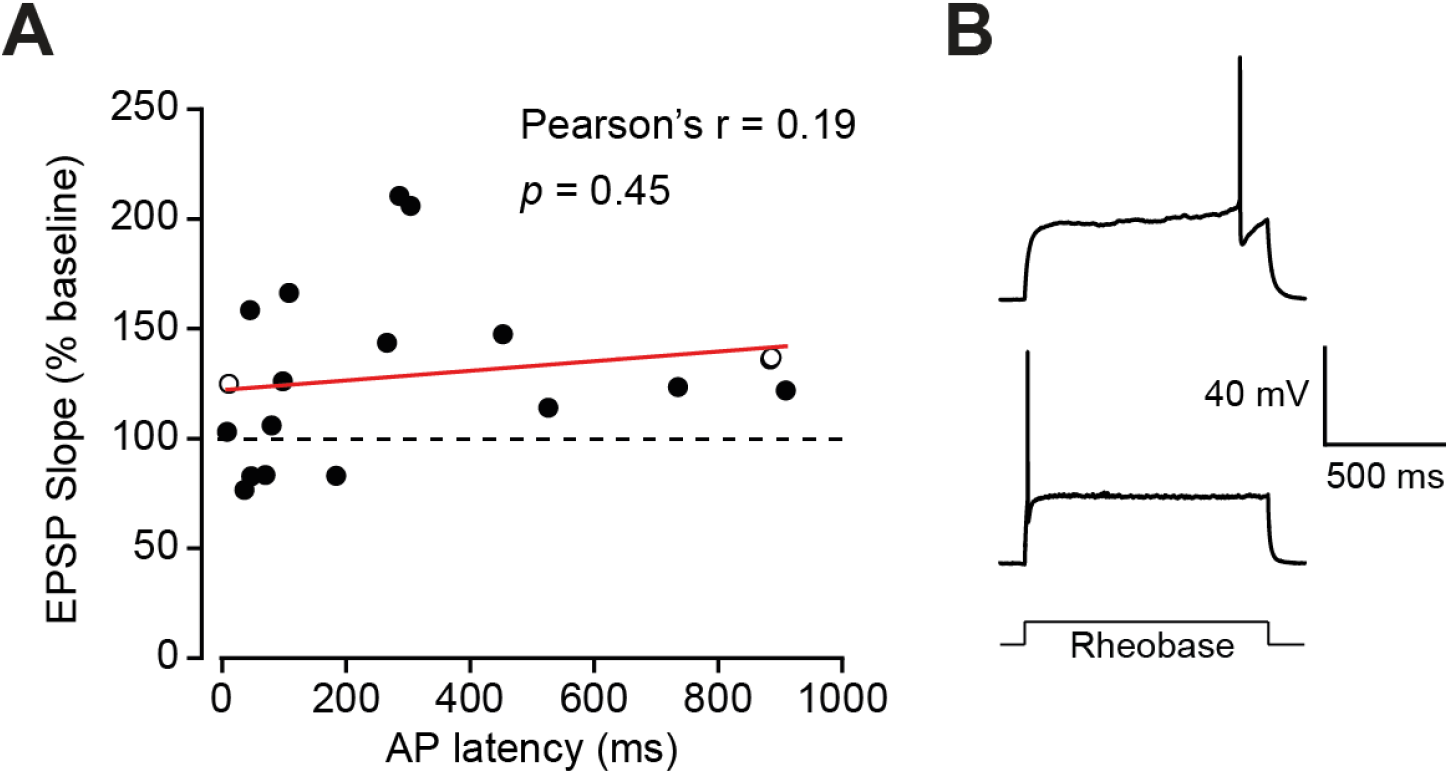
(**A**) LTP magnitude shows no obvious relationship to action potential latency at rheobase current injections. The individual cells are the same as in Figure 1*A* (n = 19). (**B**) Example traces from putative early- and late-spiking NGF cells, shown as open symbols in (A).

**Extended Data Fig. 1-2.**
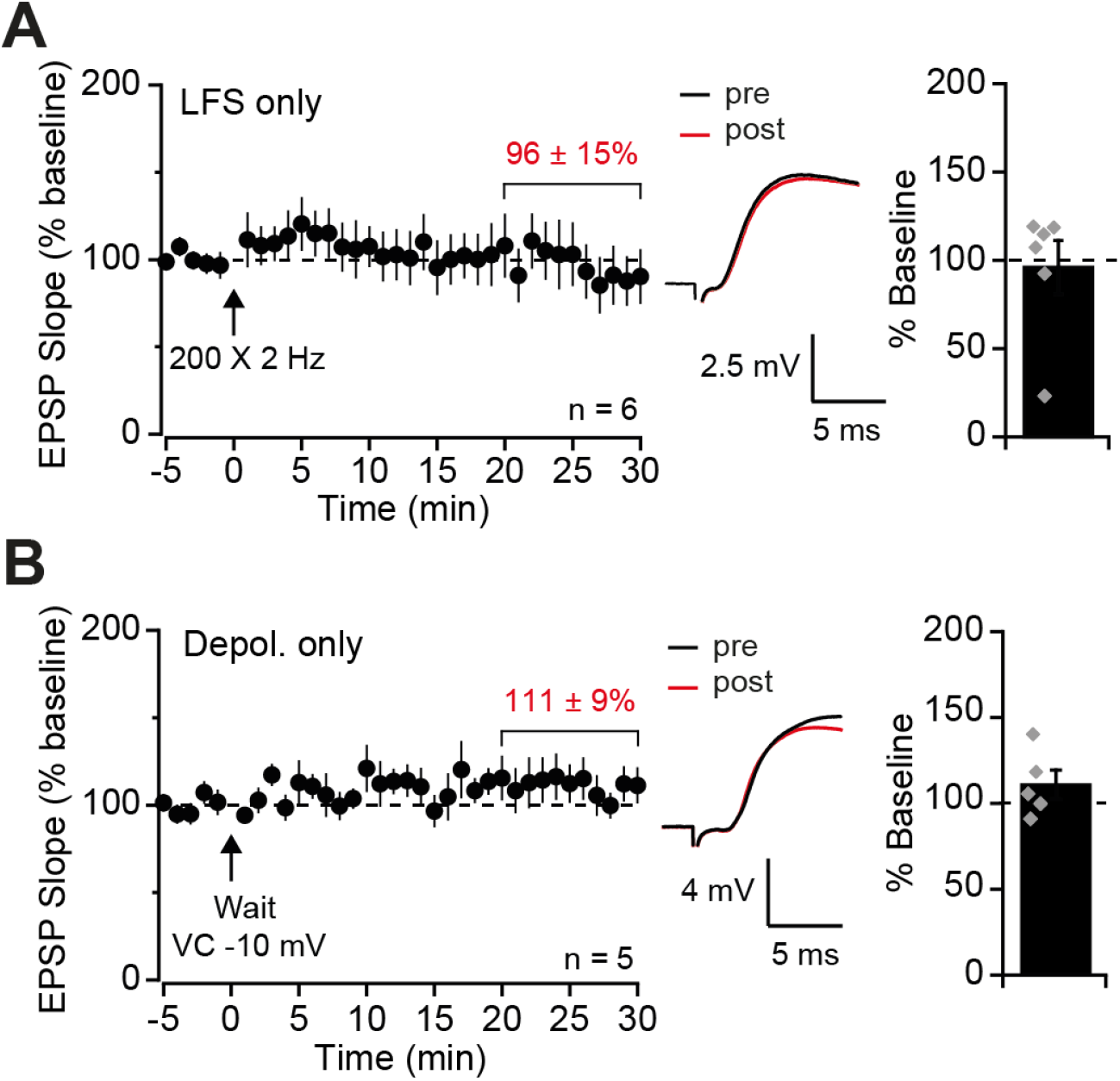
(**A**) Pooled dataset and representative traces showing no LTP in SLM interneurons following presynaptic LFS only, quantified on right. (**B**) As for (A) but for postsynaptic depolarisation only.

**Extended Data Fig. 1-3.**
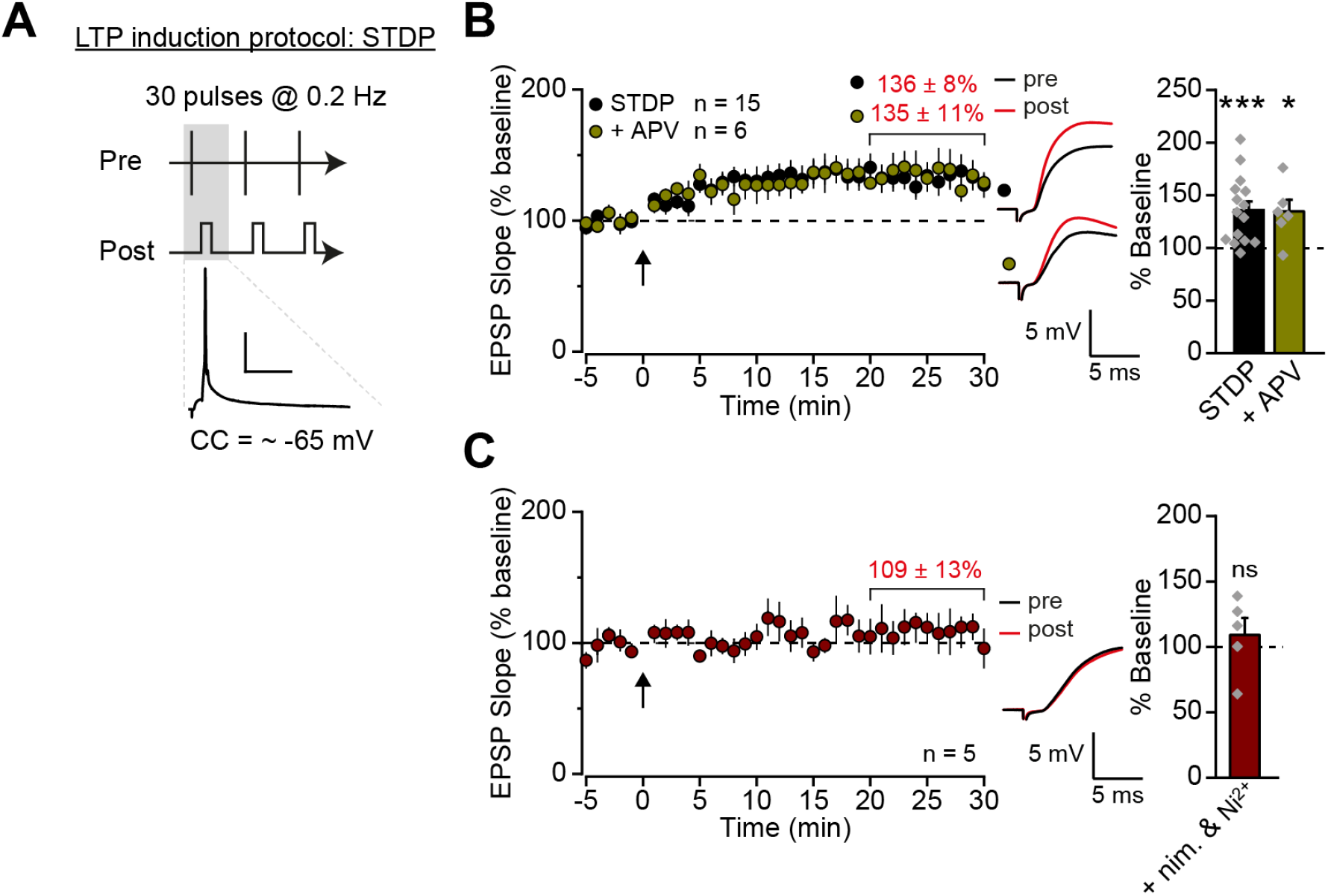
(**A**) Schematic of the spike timing-dependent plasticity (STDP) protocol, pairing 30 presynaptic stimulations with postsynaptic spikes. An example trace of a single pairing recorded from the postsynaptic cell is shown (scale: 25 mV, 40 ms). CC = current clamp. (**B**) Pooled dataset and representative traces showing that the STDP protocol induced LTP that was not NMDA receptor-dependent, quantified on right. (**C**) The STDP protocol failed to induce LTP in the presence of Ni^2+^ (100 μM) and nimodipine (10 μM; nim), quantified on right. * *P* < 0.05, *** *P* < 0.001. Error bars show SEM.

**Extended Data Fig. 2-1.**
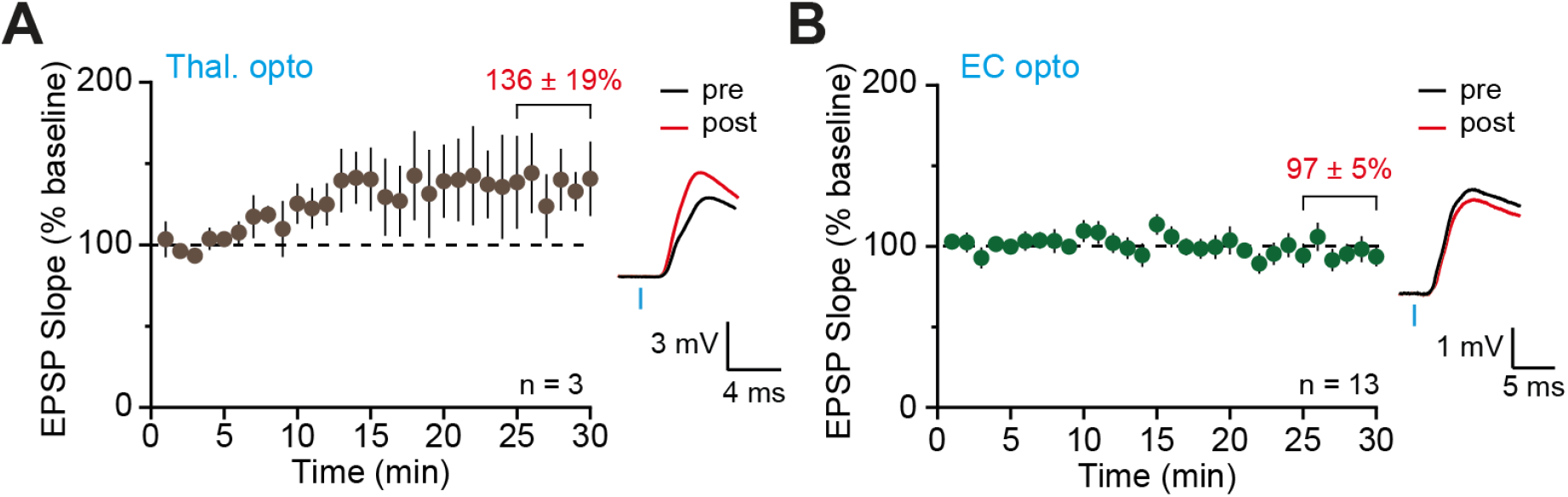
(**A**) Optogenetic stimulation of thalamic fibers evoked EPSPs in SLM interneurons that increased over a 30 minute recording period (**B**) As for (A) but showing stable recordings with optogenetic stimulation of EC fibers.

**Extended Data Fig. 5-1.**
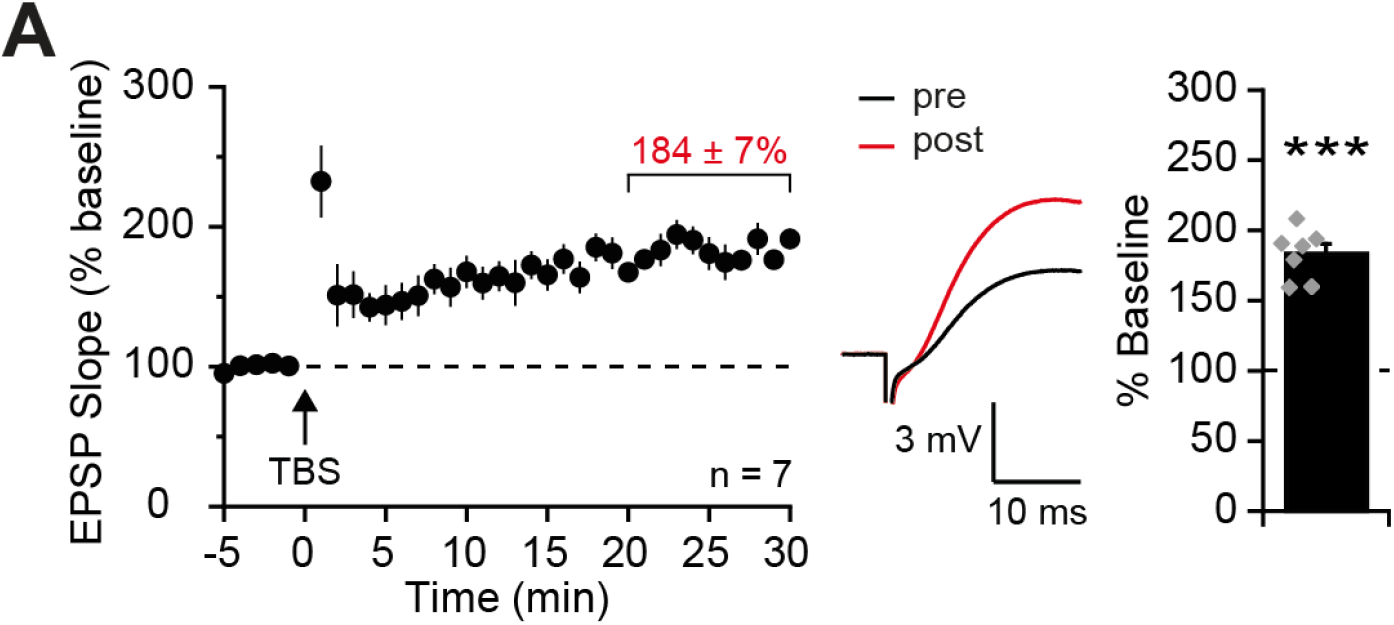
TBS of the SLM in the presence of GABA blockers induces LTP in CA1 pyramidal cells.

**Extended Data Table 3-1.** Electrophysiological properties of hippocampal NDNF+ cells

|  | *Mean ± SEM*<br>*(n = 20)* |
| --- | --- |
| *Resting potential*<br>*(mV)* | -62.1 ± 2.1 |
| *Input resistance*<br>*(MΩ)* | 221.5 ± 17.4 |
| *Time constant (τ)*<br>*(ms)* | 12.0 ± 0.5 |
| *Sag (mv)* | 1.8 ± 0.6 |
| *Sag %* | 5.1 ± 0.8 |
| *AP amplitude (mV)* | 47.8 ± 1.8 |
| *AP half-width (ms)* | 1.03 ± 0.05 |
| *AP latency (ms)* | 399.6 ± 86.8 |
| *AP threshold (mV)* | -32.6 ± 1.5 |
| *AHP amplitude (mV)* | 14.8 ± 0.9 |
| *firing frequency (Hz)* | 28.0 ± 3.1 |
| *adaptation ratio* | 0.6 ± 0.1 |

**Extended Data Table 3-2.** Properties of IPSPs in pyramidal cells evoked by optogenetic stimulation of NDNF+ cells.

|  | *Baseline*<br>(*n* = 7) | *GABA<sub>A</sub>* *<br>(*n* = 4) |
| --- | --- | --- |
| *Rise time (ms)* | 29.9 ± 2.5 | 25.3 ± 2.5 |
| *Decay (τ) (ms)\*\** | - | 44.9 ± 3.0 |
| *Amplitude (mV)* | -3.5 ± 0.7 | -5.0 ± 1.2*** |
| *Half-width (ms)* | 155.7 ± 31.1 | 72.4 ± 5.4 |

\* Isolated by application of CGP 55845 (1 μM)
\*\* Baseline responses could not be well fit by an exponential function due to their dual components.
\*\*\* Note that IPSP amplitude increased following application of CGP 55845, likely due to blockade of presynaptic GABA<sub>B</sub> receptors (Price et al., 2008).

